# Multi-Omics Characterization of Plasma and Urine Extracellular Vesicles Identifies Non-Invasive Biomarkers for IgA Nephropathy

**DOI:** 10.64898/2026.07.17.738834

**Authors:** Yun-Hsuan Lin, Tanyu Chang, I-Lin Tsai, Jayasri Parati, Chih-Chin Kao

## Abstract

**Background:** IgA nephropathy (IgAN) is increasingly recognized as a systemic immune-mediated disease characterized by aberrant IgA1 glycosylation, circulating immune complex formation, complement activation, and emerging metabolic perturbations. However, clinical diagnosis still relies on invasive renal biopsy, and non-invasive biomarkers capable of capturing both systemic immune activation and kidney-specific alterations remain lacking. Extracellular vesicles (EVs), as biologically active carriers of proteins and metabolites, provide a unique opportunity to interrogate compartment-specific molecular signatures underlying IgAN pathophysiology.

**Methods:** We performed an integrated, untargeted multi-omics analysis of plasma- and urine-derived EVs from 60 individuals (24 IgAN, 21 chronic kidney disease [CKD], and 15 controls). Differentially expressed proteins (DEPs) and metabolite features (DEFs) discriminating IgAN from CKD and controls were identified using Venn diagram analysis, followed by pathway enrichment and receiver operating characteristic (ROC) evaluation.

**Results:** Venn analysis identified 22 and 3 candidate DEPs in plasma EVs (pEVs) and urinary EVs (uEVs), respectively, revealing broader systemic proteomic alterations relative to renal EV cargo. Notably, complement and coagulation regulators, including C4b-binding protein alpha chain (C4BPA) and vitamin K–dependent protein S (PROS1), demonstrated strong discriminatory performance between IgAN and CKD (AUC = 0.826 and 0.795), suggesting EV-associated complement–coagulation crosstalk in IgAN.

Metabolomic profiling revealed 1,006 and 540 candidate DEFs in pEVs and uEVs, respectively. Enrichment analyses highlighted steroid biosynthesis and fatty acid metabolism pathways in both compartments, indicating immune–metabolic reprogramming. Three metabolite features (C27H44O, C30H50O, and C28H46O) distinguished IgAN from CKD with high accuracy (AUC = 0.942–0.877).

**Conclusions:** This study provides the first compartment-resolved, plasma- and urine-derived EV multi-omics landscape of IgAN. Our findings suggest that EV cargo reflects coordinated complement dysregulation and metabolic alterations, extending current understanding of IgAN beyond glomerular immune complex deposition. These EV-associated proteins and metabolites offer a mechanistically informed framework for non-invasive biomarker development and for exploring immune–metabolic pathways involved in IgAN progression.

## Introduction

IgA nephropathy (IgAN) is the most common form of primary glomerulonephritis (GN) worldwide and a leading cause of chronic kidney disease[1–6]. The disease is characterized by the deposition of abnormally glycosylated galactose-deficient IgA1 (Gd-IgA1) in the glomerular mesangium, often accompanied by IgG, IgM, and complement C3, resulting in mesangial cell proliferation and excessive extracellular matrix production[7].

The global prevalence of IgAN varies by region, with a particularly high incidence in the Asia-Pacific region. However, as diagnosis requires an invasive kidney biopsy, the actual disease prevalence is likely underestimated, especially in patients with mild or asymptomatic conditions[7, 8]. IgAN typically affects adults between their twenties and thirties, with an estimated incidence of at least 2.5 cases per 100,000 individuals. Clinically, IgAN may present with painless hematuria, proteinuria, back pain, edema, and progressive renal dysfunction, while some patients remain asymptomatic for years[9]. Notably, approximately 30% of patients progress to end-stage renal disease (ESRD) within 10 to 20 years of onset, emphasizing the need for early diagnosis and intervention[1, 3, 10, 11].

Although the exact etiology of IgAN remains unclear, the “multi-hit hypothesis” is the most widely accepted model explaining its pathogenesis[5]. This hypothesis describes four major steps: 1. Overproduction of Gd-IgA1 with aberrant O-glycosylation in the hinge region. 2. Recognition of Gd-IgA1 by autoantibodies (IgG or IgM), forming immune complexes. 3. Deposition of these complexes in glomerular mesangial cells. 4. Activation of inflammatory pathways, leading to mesangial proliferation, complement activation, glomerulosclerosis, and tubular atrophy[1, 3–5, 11–15].

Despite significant advances in understanding the disease mechanism, clinical diagnosis remains challenging. Symptoms are nonspecific[4], and renal biopsy—with immunofluorescence detection of mesangial IgA, IgG, IgM, and C3 deposits— remains the only definitive diagnostic method. The Oxford MEST-C classification is used for grading histopathological severity. However, due to the invasiveness of biopsy, early or mild IgAN cases are often undiagnosed. Currently, no reliable non-invasive biomarkers are available for IgAN diagnosis, underscoring the need for novel diagnostic tools.

Extracellular vesicles (EVs) have recently emerged as a crucial mode of intercellular communication, mediating the transfer of proteins, lipids, and nucleic acids between cells. Initially regarded as cellular waste disposal vehicles, EVs are now recognized as selective carriers of biologically active molecules that regulate cellular functions in both physiological and pathological states[16].

EVs are lipid bilayer-enclosed vesicles released by almost all cell types and are broadly classified into exosomes, microvesicles, apoptotic bodies, and migrasomes, depending on their size and biogenesis pathway[17, 18]. They circulate in various body fluids—including plasma, urine, cerebrospinal fluid, and amniotic fluid—and reflect the molecular phenotype of their parent cells.[19, 20] This stability and accessibility make EVs promising candidates for non-invasive biomarker discovery in a wide range of diseases[21].

EVs are increasingly recognized as valuable diagnostic tools because they mirror the physiological and pathological status of their originating cells. For instance, specific EV-associated proteins such as Fetuin-A have been identified as biomarkers for acute kidney injury and cancers[4]. Plasma EVs (pEVs) often originate from immune cells under inflammatory or allergic conditions[22], while urinary EVs (uEVs) carry signatures of renal and urogenital health[23].

In kidney diseases, uEVs can reflect glomerular and podocyte injury, offering a more comprehensive view of disease mechanisms compared to conventional urine or plasma biomarkers[4]. Among EV subtypes, small EVs (30–150 nm) have shown particularly high potential for disease monitoring[24].

Previous studies have explored EVs in IgAN through proteomic analyses. For example, Pyong-Gon Moon et al. compared urinary EV proteins from IgAN and thin basement membrane nephropathy patients[4], while Negin Farzamikia et al. analyzed EV protease activity in IgAN versus healthy controls[25]. However, no study to date has simultaneously analyzed both plasma and urinary EVs while including chronic kidney disease (CKD) as a comparative disease group. Such integrative analysis could enable the identification of early, non-invasive biomarkers specific to IgAN, addressing a critical gap in current diagnostic approaches.

In this study, extracellular vesicles from purified blood and urine clinical samples will be subjected to proteomic and metabolomic analysis, comparing protein and metabolite expression levels between IgA nephropathy (IgAN) patients, chronic kidney disease (CKD) patients, and control groups. The aim is to identify potential proteins and metabolites to aid in the development of early diagnostic biomarkers.

## Methods

### Study design and patient enrollment

This was a comparison study evaluating plasma and urinary EV proteomic and metabolomic profiles in patients with IgAN, CKD, and controls. The overall study design is illustrated **(Figure 1)**. Patients with IgAN and CKD were recruited from Taipei Medical University Hospital (Taipei, Taiwan) between April 10, 2013, and January 7, 2022. The study was approved by the Taipei Medical University Joint Institutional Review Board (IRB No. N201704064), and informed consent was obtained from all participants. Sixty subjects were enrolled in this study, including 24 patients with IgAN, 21 with chronic kidney disease, and 15 controls. Baseline demographic and clinical data were collected. Differentially expressed proteins (DEPs) and differentially expressed features (DEFs) that both discriminate IgAN from CKD and Control were identified as candidate biomarkers through Venn diagram analysis.

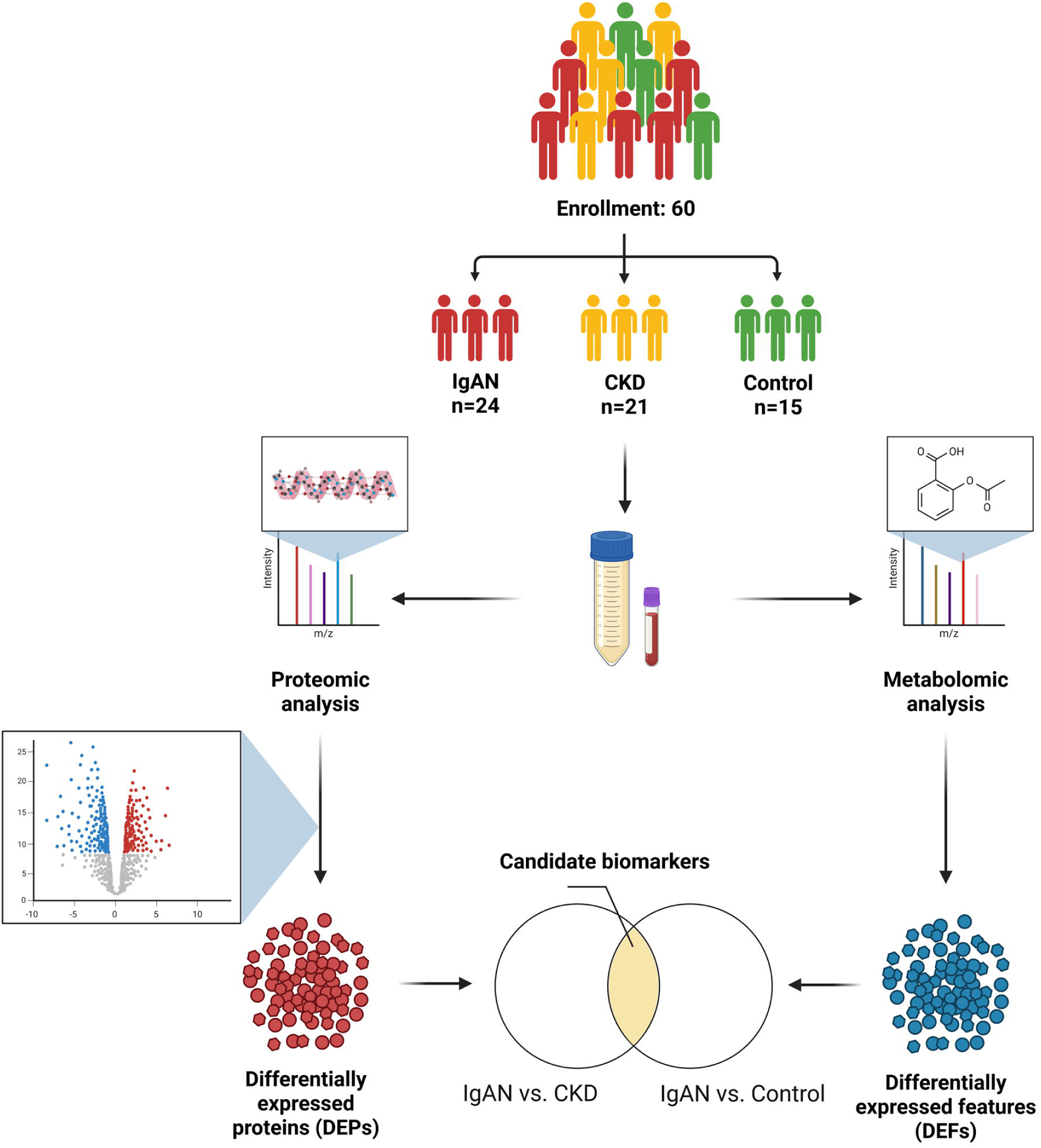

### EV isolation and characterization

Plasma and urinary EVs were isolated as outlined in **Figure 1A**. Briefly, after sample collection, 15 mL of urine was additionally concentrated to 500 µL using Amicon Ultra-15 centrifugal filters (100 kDa cutoff; Merck Millipore, Darmstadt, Germany). EVs were purified using the SmartSEC™ HT EV Isolation System (System Biosciences, Palo Alto, CA, USA), following the manufacturer’s instructions[26]. To confirm EV isolation, the following validation techniques were employed: first, western blotting for exosomal markers CD63 and CD9, as well as the negative EV marker apolipoprotein A-I (APOA-1) (System Biosciences, Palo Alto, CA, USA)[27], with 20 μg of protein per sample was performed. Second, nanoparticle tracking analysis (NTA) using the NanoSight NS300 (Malvern Panalytical, UK) to assess particle size and concentration was done. Finally, transmission electron microscopy (TEM) using a Hitachi HT-7700 (Hitachi, Japan) to examine EV morphology was performed.

### Sample preparation for proteomic and metabolomic analysis

Proteins were precipitated with ice-cold acetone (Honeywell, Charlotte, NC, USA) at a volume four times that of each sample, and 50 µg of protein was processed per sample. Pellets were resuspended in 100 µL of 6 M urea (Nihon Shiyaku Industries, LTD, Japan), reduced with 2 µL of 550 mM dithiothreitol (DTT; Sigma-Aldrich, St. Louis, MO, USA), alkylated with 4 µL of 450 mM iodoacetamide (IAA; Sigma-Aldrich), and digested with 5 µL of trypsin (0.2 µg/µL; Promega, Madison, WI, USA). Solid-phase extraction (SPE) was performed using Sep-Pak® Vac 1cc (50mg) C18 cartridges (Waters, Milford, MA, USA) and a SpeedVac concentrator to purify and concentrate peptides. Meanwhile, the supernatant was collected for metabolomic analysis. Like proteomic analysis, metabolites were concentrated using a SpeedVac concentrator. The dried extracts were reconstituted in 50% methanol for LC–MS/MS analysis.

### Liquid chromatography-mass spectrometry (LC-MS)

For proteomic analysis, MS and gradient settings optimized in our previous work[28] were applied. Analyses were performed on an Orbitrap Fusion Lumos Tribrid quadrupole-ion trap-Orbitrap mass spectrometer (Thermo Fisher Scientific, CA, USA) coupled with an Ultimate 3000 NanoLC system (Thermo Fisher Scientific, Bremen, Germany). Peptides were separated on a C18 Acclaim PepMap NanoLC column (75 μm ID × 25 cm). A segmented gradient (2%–40% acetonitrile in 0.1% formic acid) was applied over 40 minutes at a flow rate of 300 nL/min.

A full MS scan was conducted, followed by high-energy collision-activated dissociation (HCD)-MS/MS analysis of the most intense ions within 3 s. Data were externally calibrated to maintain a mass accuracy of <5 ppm. MS1 scans were acquired at a resolution of 120,000 (m/z 200), followed by HCD-MS/MS scans (resolution: 15,000) with dynamic exclusion (60 s), a 1.4 Da isolation window, and a normalized collision energy of 32%. The raw data were deposited in the ProteomeXchange Consortium via PRIDE (dataset identifier: PXD074396).

For metabolomic analysis, data acquisition was performed on a SYNAPT XS mass spectrometer (Waters, MA, USA) equipped with an ACQUITY UPLC system (Waters, Milford, MA, USA). Metabolites were separated on a Poroshell 120 EC-C18 column (1.9 μm, 2.1 × 100 mm). MS1 scanning ranges from 50 Da to 1200 Da, with a capillary voltage maintained at 2.0 kV, a source temperature of 150 °C, a desolvation temperature of 400 °C, a cone voltage of 40 V, and a cone gas flow of 50 L/h. The cone gas flow is 50 L/hr. The survey scan starts at a mass of 50 Da and ends at 1000 Da, with an intensity threshold of 10,000 and the detection of up to 10 components.

### Data Processing and Bioinformatics Analysis

Protein identification and label-free quantification were performed using MaxQuant (v2.4.7.0) with a reviewed human FASTA database (UniProtKB Taxonomy ID: 9606; downloaded September 12, 2023). Data were filtered and normalized in Perseus (v2.0.11). Proteins with missing values in more than 60% of the samples were excluded from the analysis. Downstream analyses—hierarchical clustering, sPLS-DA, and volcano plots—were conducted in MetaboAnalyst 6.0. DEPs are defined as proteins with a Wilcoxon test p-value < 0.05 and fold change > 1.3.

Metabolite identification and label-free quantification were conducted using Progenesis QI with the Human Metabolome Database (Metabolite Structures, Version 5.0, downloaded August 20, 2025). Features with missing values in more than 60% of the samples were excluded from further analysis. Downstream analyses, hierarchical clustering, sPLS-DA, and volcano plots were also conducted in MetaboAnalyst 6.0. DEFs were defined as features with a Wilcoxon test p-value < 0.05 and fold change >1.3. Gene enrichment analysis of the DEPs was performed using FunRich (v3.1.3).

## Results

### Patients’ demographics and outcomes

A total of 60 subjects were enrolled in the study, including 24 patients in the IgAN group, 21 in the CKD group, and 15 in the Control group. Patient characteristics are summarized in **Table 1**. The mean age was 55.6 years, and 28 (46.7%) were male. Baseline characteristics were generally comparable between the groups, except for differences in renal function, which were highest in the Control group and lowest in the CKD group.

**Table 1.**
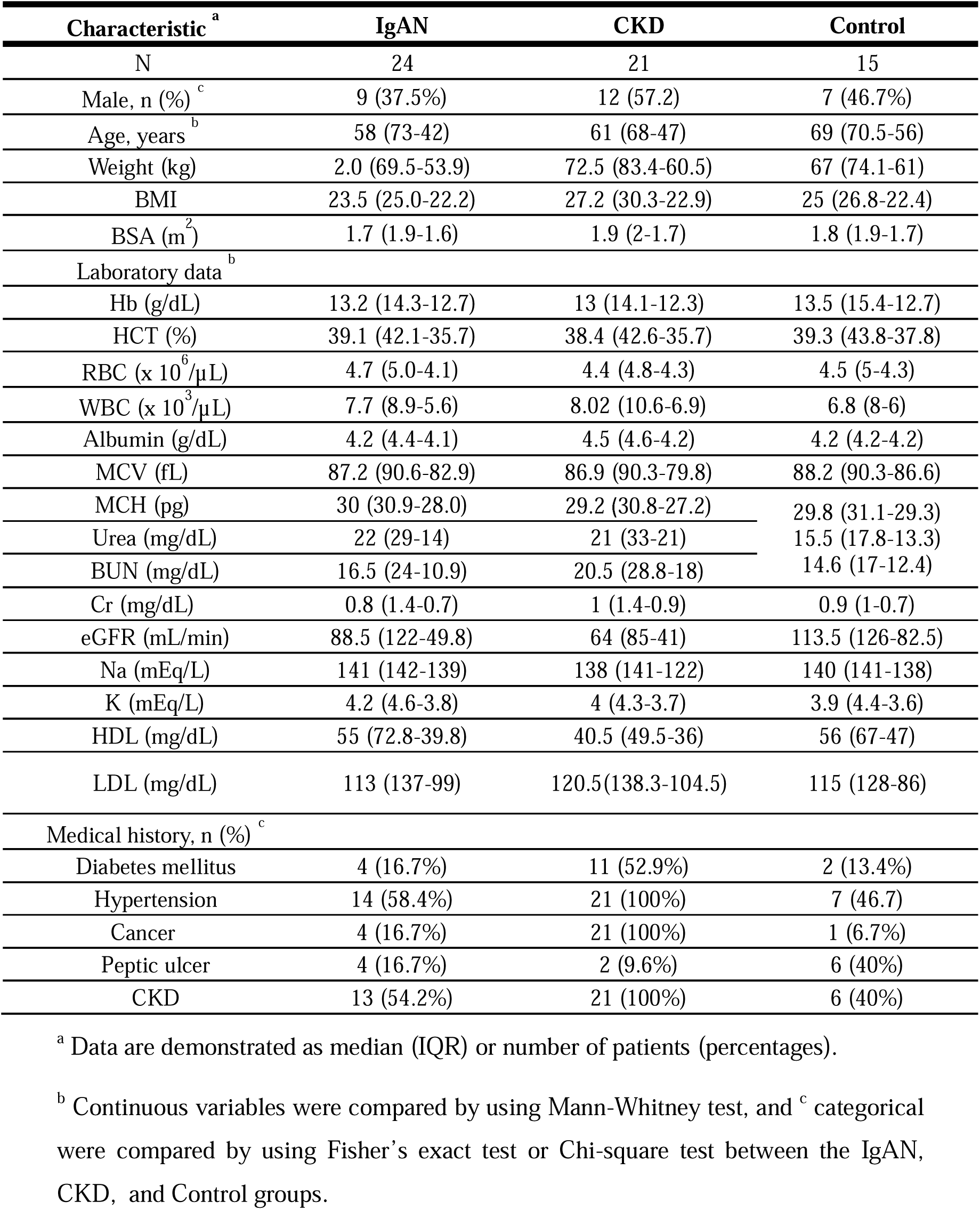
Baseline characteristics of IgAN, CKD patients, and Controls.

| Characteristic <sup>a</sup> | IgAN | CKD | Control |
| --- | --- | --- | --- |
| N | 24 | 21 | 15 |
| Male, n (%) <sup>c</sup> | 9 (37.5%) | 12 (57.2) | 7 (46.7%) |
| Age, years <sup>b</sup> | 58 (73-42) | 61 (68-47) | 69 (70.5-56) |
| Weight (kg) | 2.0 (69.5-53.9) | 72.5 (83.4-60.5) | 67 (74.1-61) |
| BMI | 23.5 (25.0-22.2) | 27.2 (30.3-22.9) | 25 (26.8-22.4) |
| BSA (m<sup>2</sup>) | 1.7 (1.9-1.6) | 1.9 (2-1.7) | 1.8 (1.9-1.7) |
| Laboratory data <sup>b</sup> |  |  |  |
| Hb (g/dL) | 13.2 (14.3-12.7) | 13 (14.1-12.3) | 13.5 (15.4-12.7) |
| HCT (%) | 39.1 (42.1-35.7) | 38.4 (42.6-35.7) | 39.3 (43.8-37.8) |
| RBC (x 10<sup>6</sup>/μL) | 4.7 (5.0-4.1) | 4.4 (4.8-4.3) | 4.5 (5-4.3) |
| WBC (x 10<sup>3</sup>/μL) | 7.7 (8.9-5.6) | 8.02 (10.6-6.9) | 6.8 (8-6) |
| Albumin (g/dL) | 4.2 (4.4-4.1) | 4.5 (4.6-4.2) | 4.2 (4.2-4.2) |
| MCV (fL) | 87.2 (90.6-82.9) | 86.9 (90.3-79.8) | 88.2 (90.3-86.6) |
| MCH (pg) | 30 (30.9-28.0) | 29.2 (30.8-27.2) | 29.8 (31.1-29.3) |
| Urea (mg/dL) | 22 (29-14) | 21 (33-21) | 15.5 (17.8-13.3) |
| BUN (mg/dL) | 16.5 (24-10.9) | 20.5 (28.8-18) | 14.6 (17-12.4) |
| Cr (mg/dL) | 0.8 (1.4-0.7) | 1 (1.4-0.9) | 0.9 (1-0.7) |
| eGFR (mL/min) | 88.5 (122-49.8) | 64 (85-41) | 113.5 (126-82.5) |
| Na (mEq/L) | 141 (142-139) | 138 (141-122) | 140 (141-138) |
| K (mEq/L) | 4.2 (4.6-3.8) | 4 (4.3-3.7) | 3.9 (4.4-3.6) |
| HDL (mg/dL) | 55 (72.8-39.8) | 40.5 (49.5-36) | 56 (67-47) |
| LDL (mg/dL) | 113 (137-99) | 120.5(138.3-104.5) | 115 (128-86) |
| Medical history, n (%) <sup>c</sup> |  |  |  |
| Diabetes mellitus | 4 (16.7%) | 11 (52.9%) | 2 (13.4%) |
| Hypertension | 14 (58.4%) | 21 (100%) | 7 (46.7) |
| Cancer | 4 (16.7%) | 21 (100%) | 1 (6.7%) |
| Peptic ulcer | 4 (16.7%) | 2 (9.6%) | 6 (40%) |
| CKD | 13 (54.2%) | 21 (100%) | 6 (40%) |

<sup>a</sup> Data are demonstrated as median (IQR) or number of patients (percentages).
<sup>b</sup> Continuous variables were compared by using Mann-Whitney test, and <sup>c</sup> categorical were compared by using Fisher's exact test or Chi-square test between the IgAN, CKD, and Control groups.

### Characterization of urinary extracellular vesicles

Isolated urinary EVs displayed a heterogeneous size distribution and exhibited the characteristic cup-shaped morphology, consistent with previous reports [11] **(Figure 2B, C)**. Western blot analysis confirmed the presence of exosomal markers CD63 and CD9 in both groups **(Figure 2D)**.

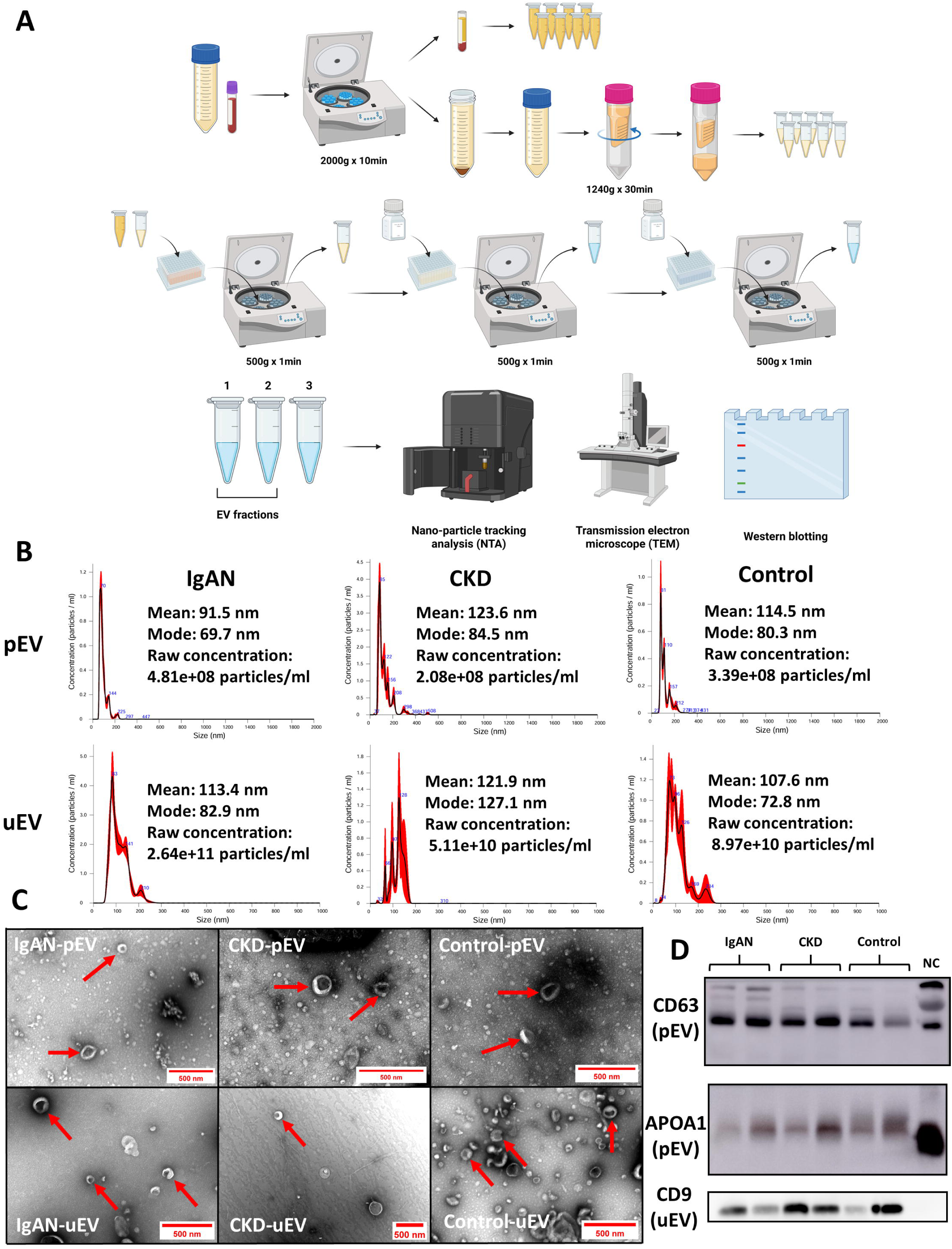

### Discovery of candidate pEV and uEV biomarkers

Heatmaps of the proteomic data showed clear groupwise clustering in both the pEV and uEV batches, with most proteins up-regulated in the IgAN group relative to CKD and controls **(Figure 3A, B)**. sPLS-DA score plots likewise demonstrated clear separation among IgAN, CKD, and controls in both batches, with component 1 explaining 20.6% and 14.6% of the variance, respectively **(Figure 3C, D)**. Volcano plots identified 95 DEPs in the pEV batch (35 in the IgAN–vs–CKD dataset; 60 in the IgAN–vs–Control dataset) and 49 DEPs in the uEV batch (22 in IgAN–vs–CKD; 27 in IgAN–vs–Control) **(Figure S1A–D)**.

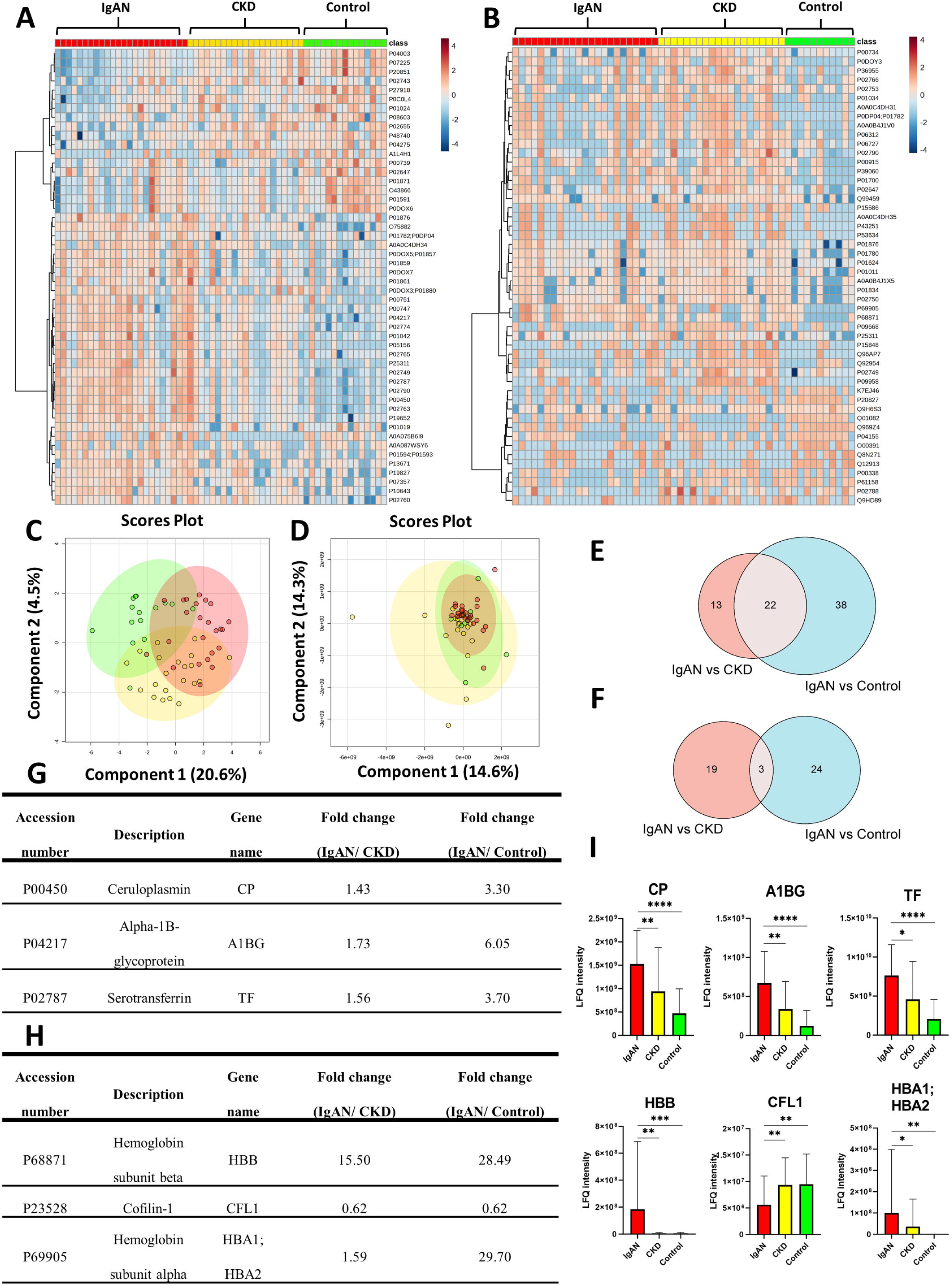

Venn diagrams yielded 22 and 3 candidate DEPs in the pEV and uEV batches, respectively **(Figure 3E, F)**. Since most candidates were complement-related proteins, we prioritized three biomarkers that are more directly linked to disease pathogenesis beyond complement activation. The three candidate DEPs and their expression patterns across IgAN, CKD, and controls are shown in **Figure 3G–I**.

Heatmaps of the metabolomic data also showed clear groupwise clustering **(Figure 4A, B; S2A, B)**. sPLS-DA plots demonstrated clear separation among IgAN, CKD, and controls across all datasets, with component 1 explaining 8.2%–12.7% of the variance **(Figure 4C, D; S2C, D)**. In positive mode, volcano plots identified 2,697 DEFs in the pEV batch (1,209 in IgAN–vs–CKD; 1,488 in IgAN–vs–Control) and 1,934 DEFs in the uEV batch (889 in IgAN–vs–CKD; 1,045 in IgAN–vs–Control) **(Figure S3A–D)**. In negative mode, 853 DEFs were found in the pEV batch (400 in IgAN–vs–CKD; 453 in IgAN–vs–Control) and 832 in the uEV batch (408 in IgAN– vs–CKD; 424 in IgAN–vs–Control) **(Figure S3E–H)**.

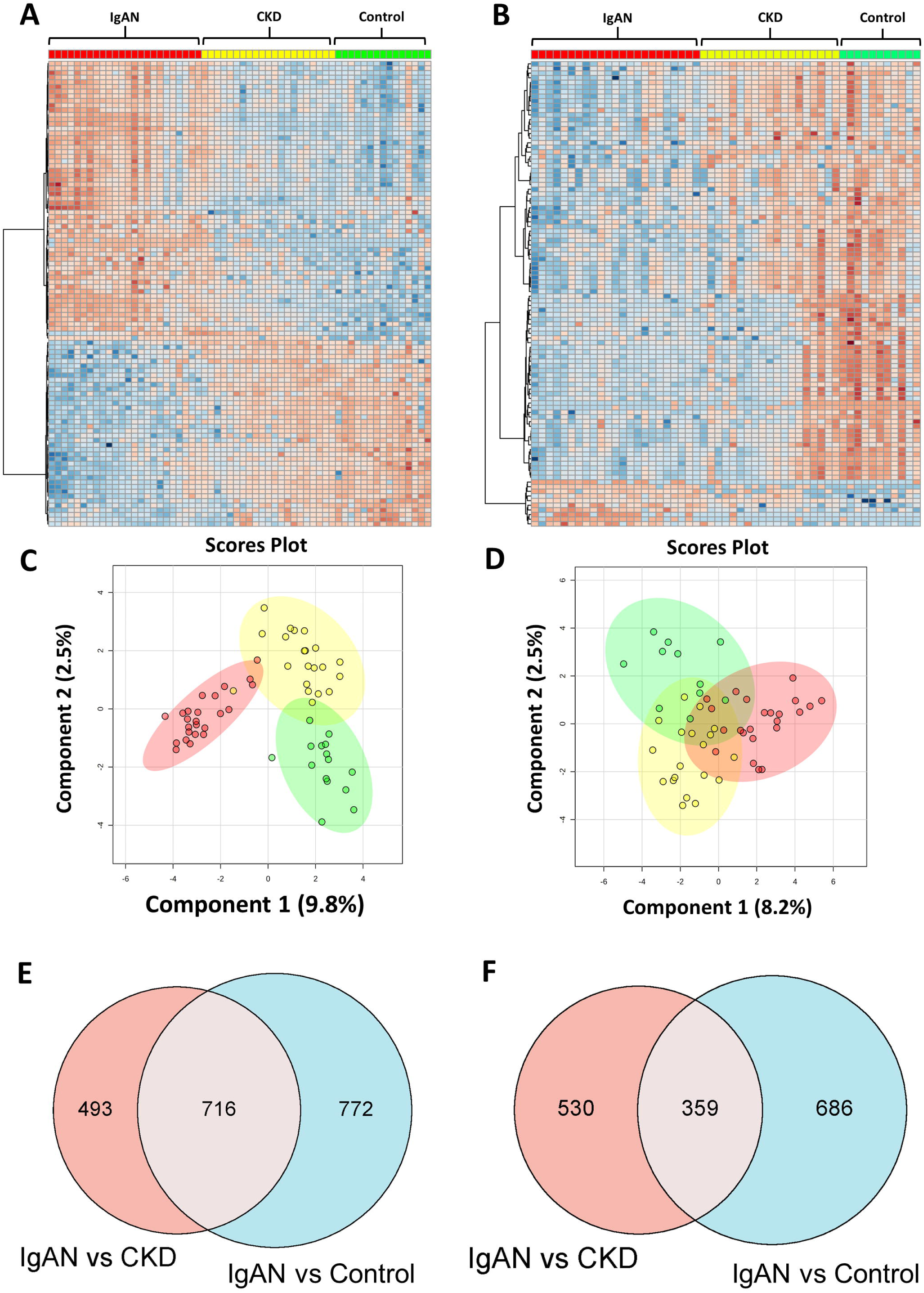

Venn analyses yielded 1,006 (716 positive; 290 negative) and 540 (359 positive; 181 negative) candidate DEFs that discriminated IgAN from both CKD and controls in the pEV and uEV batches, respectively **(Figure 4E, F; S2E, F)**. Because authentic reference standards were unavailable, compound IDs remained putative, which were difficult to interpret; accordingly, pathway analyses were performed to prioritize desirable metabolites for subsequent validation of metabolomic DEFs. Among enriched pathways, Steroid biosynthesis, Valine, leucine, and isoleucine biosynthesis, and Linoleic acid metabolism were most prominent in the pEV batch, whereas Linoleic acid metabolism, Arachidonic acid metabolism, and Steroid hormone biosynthesis were most enriched in the uEV batch **(Figure 5A, B)**. Based on a literature review, we prioritized steroid biosynthesis because it was enriched in both datasets and is strongly supported by prior studies.

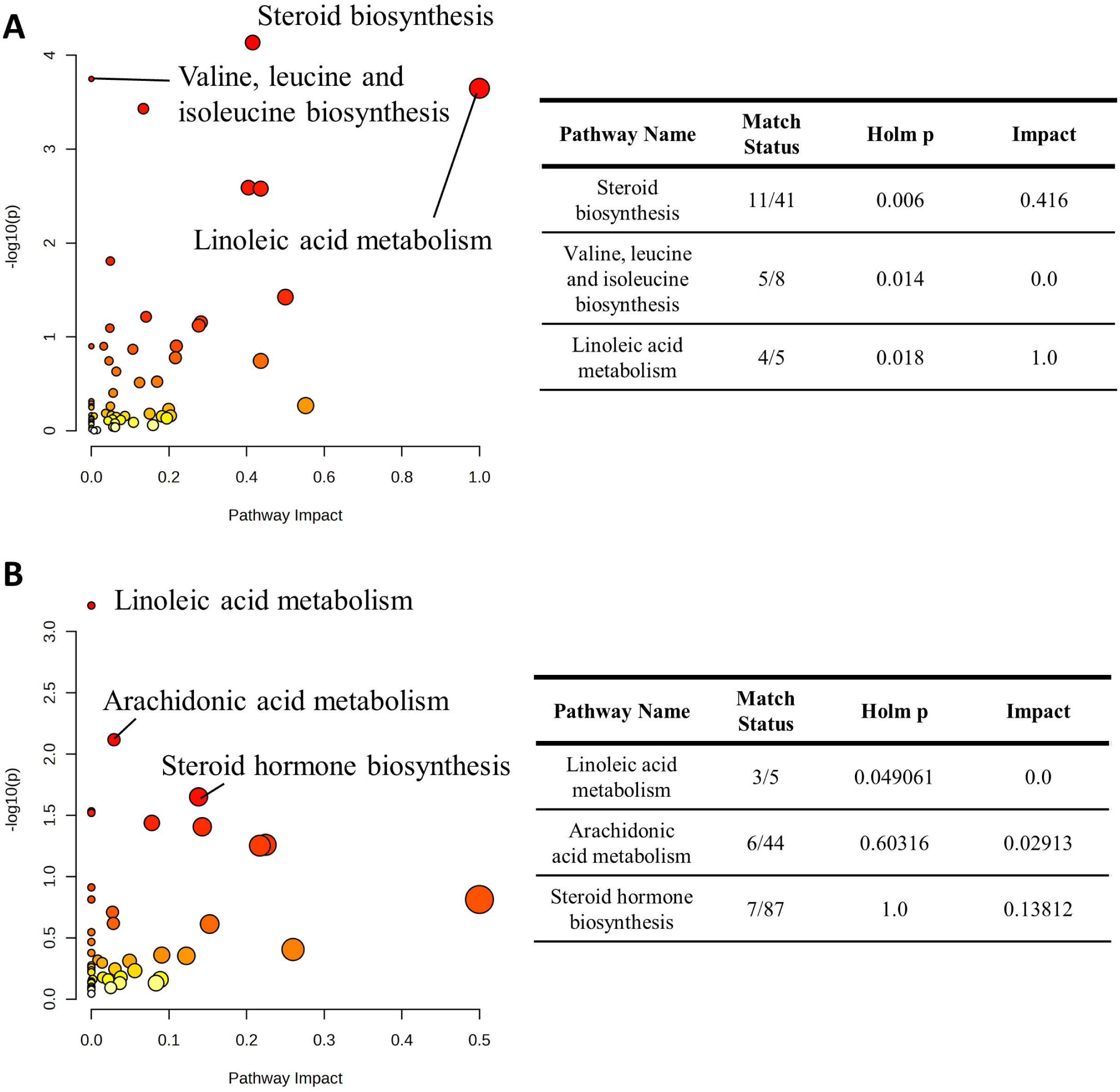

### Validation of Candidate Biomarkers for IgAN

For proteomics, receiver operating characteristic (ROC) analysis in the pEV batch showed that C4b-binding protein alpha chain (C4BPA) distinguished IgAN from CKD with an AUC of 0.826 (95% CI, 0.692–0.928). Several other DEPs also demonstrated diagnostic utility: vitamin K–dependent protein S (PROS1), AUC 0.806 (95% CI, 0.683–0.910); C4b-binding protein beta chain (C4BPB), AUC 0.795 (95% CI, 0.664–0.897); alpha-1-acid glycoprotein 1 (ORM1), AUC 0.770 (95% CI, 0.610–0.894); and vitamin D–binding protein (GC), AUC 0.751 (95% CI, 0.581–0.883). In the uEV batch, hemoglobin subunit beta (HBB) also showed good diagnostic performance, with an AUC of 0.770 (95% CI, 0.615–0.902) **(Figure 6)**. Notably, ROC analyses for each candidate DEP exhibited strong discrimination of IgAN versus controls **(Figure S4)**.

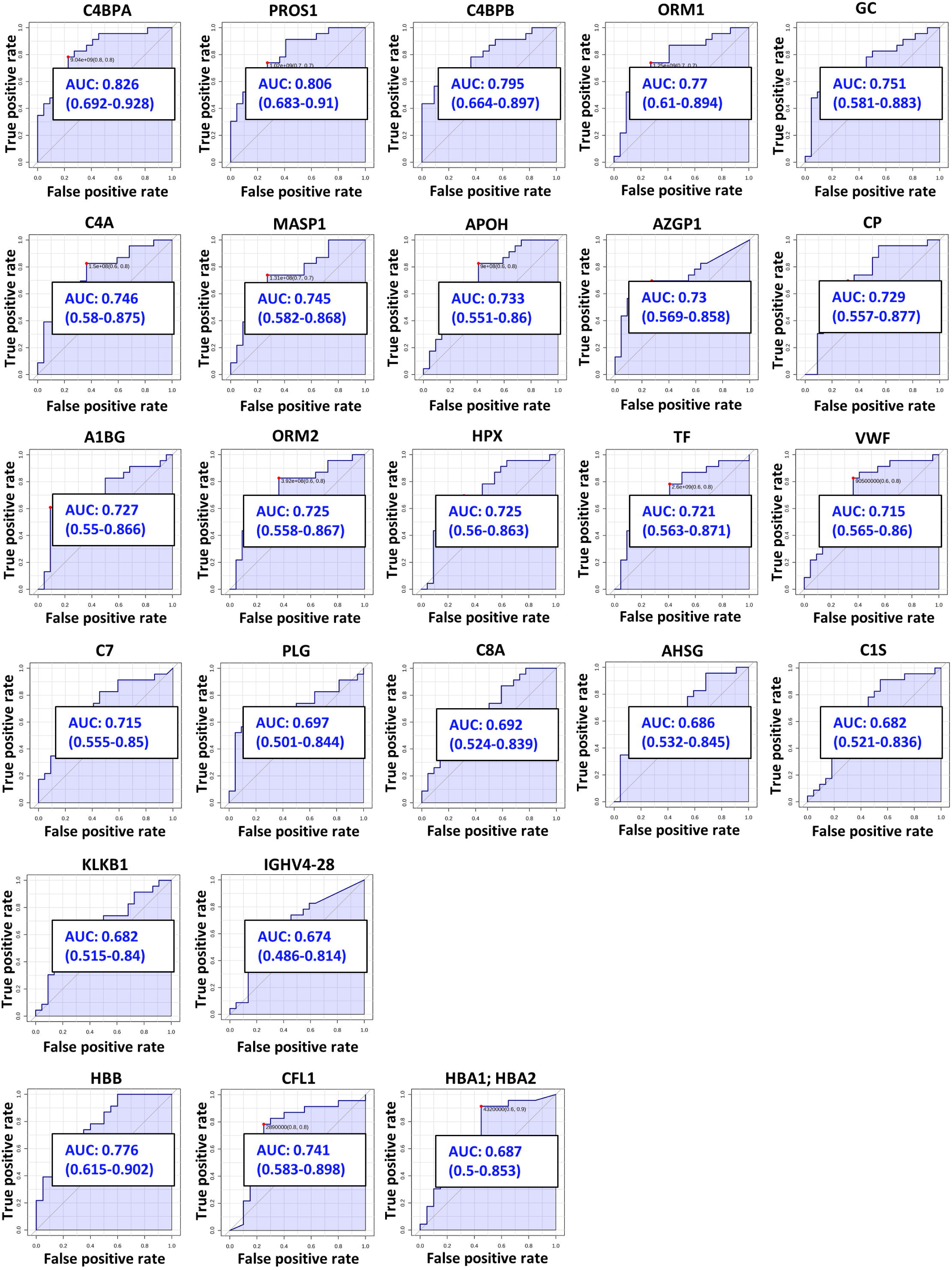

For metabolomics, ROC analysis in the pEV batch showed that features with formulas C27H44O, C30H50O, and C28H46O discriminated IgAN from CKD with AUCs of 0.942, 0.915, and 0.877, respectively. By contrast, in the uEV batch, all features exhibited relatively weak discrimination of IgAN versus CKD **(Figure 7)**.

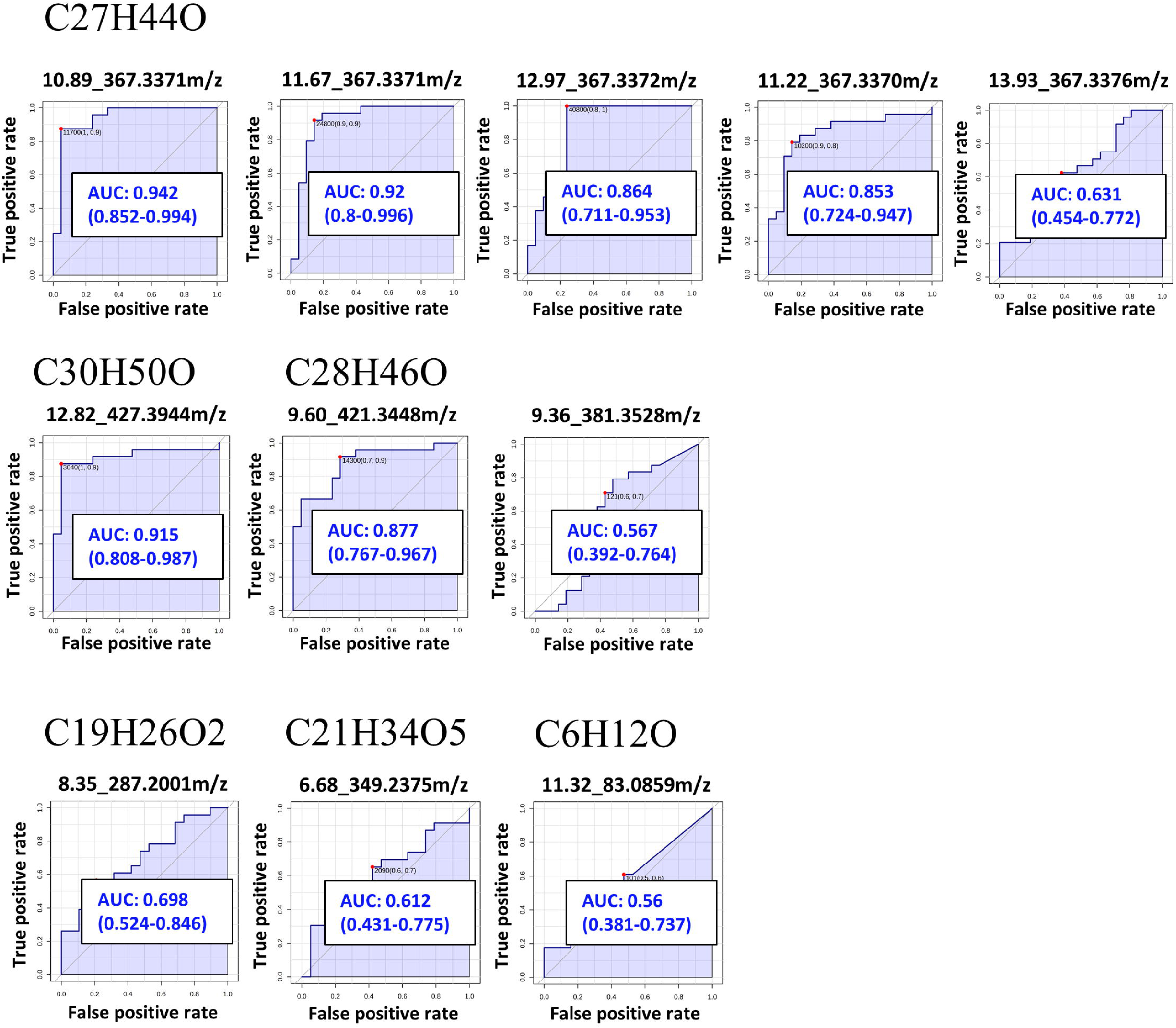

## Discussion

The present study is, to our knowledge, the first to perform a non-targeted, bottom-up multi-omics analysis integrating both plasma- and urine-derived extracellular vesicles (EVs) from patients with IgA nephropathy (IgAN). Previous EV-based studies in IgAN have largely focused on genomic or transcriptomic approaches[29–33], while proteomic investigations using EVs for biomarker discovery remain limited. For instance, Moon et al. conducted proteomic profiling of urinary exosomes from patients with early IgAN and thin basement membrane nephropathy (TBMN), identifying candidate biomarkers capable of discriminating between the two conditions[4]. In addition, although a few previous multi-omics studies identified diagnostic markers for IgAN[10, 34], the biological sources were restricted to plasma and urine. By contrast, our study simultaneously interrogates plasma EVs and urine EVs, thereby providing a more comprehensive and systemic view of EV-associated molecular alterations in IgAN.

Our proteomic findings are supported by several previous studies. Mucha *et al.* reported that zinc-alpha-2-glycoprotein, ceruloplasmin, and vitamin D-binding protein (GC) were markedly up-regulated in IgAN compared with healthy controls[35]. Consistent with these results, all three proteins were also significantly increased in the IgAN group relative to both the CKD and Control groups in our cohort. Similarly, Kalantari *et al.* employed complementary nanoflow LC-MS/MS and GeLC-MS/MS approaches and identified significant up-regulation of GC and hemopexin (HPX) in IgAN[36]. These proteins were likewise elevated in our dataset. Mohammadi Majd *et al.* further constructed a diagnostic biomarker panel based on urinary proteomic profiles, identifying transferrin (TF) as the most prominently up-regulated and diagnostically informative marker[37]; notably, TF was also increased in our study, further supporting its diagnostic relevance in IgAN.

In addition to these established markers, hemoglobin-related proteins and cytoskeleton-associated proteins identified in our EV datasets are also supported by prior evidence. Li *et al.* reported significant up-regulation of hemoglobin subunit beta (HBB) in renal biopsy transcriptomic datasets from patients with IgAN, consistently observed in both glomerular and tubulointerstitial compartments[38]. Rudnicki *et al.* developed a urinary peptide–based algorithm for predicting IgAN progression, in which both hemoglobin subunit beta and hemoglobin subunit alpha were included[39]. Moreover, Zhao *et al.* demonstrated that urinary cofilin-1, a podocyte-related actin-binding protein, was significantly elevated in IgAN patients and independently associated with baseline renal function decline and adverse renal outcomes[40]. Collectively, these findings support the biological plausibility of HBB and cofilin-1 as EV-associated markers reflecting renal injury and cytoskeletal remodeling in IgAN.

Accumulating evidence has demonstrated that dysregulation of the complement system plays a central role in the pathogenesis of IgA nephropathy[10, 11, 35]. In line with this concept, our proteomic analysis revealed that a substantial proportion of candidate DEPs were enriched in complement activation–related pathways, further supporting the involvement of complement-mediated immune mechanisms in IgAN and highlighting the relevance of EVs as carriers of complement-associated signals.

In the metabolomic analysis, two major pathways were consistently enriched in both plasma EV and urine EV datasets. Linoleic acid metabolism has previously been reported to be significantly disrupted in IgAN, as demonstrated by an untargeted metabolomic analysis of intestinal metabolites. Wu *et al.* showed that key epoxy metabolites within the linoleic acid pathway were markedly reduced in IgAN patients, suggesting suppression of linoleic acid metabolic activity and a concomitant decrease in anti-inflammatory lipid mediators[41]. Such metabolic alterations may contribute to immune dysregulation and intestinal barrier dysfunction in IgAN. In addition, candidate metabolites in our EV datasets were enriched in steroid biosynthesis and steroid hormone biosynthesis pathways. Although corticosteroid therapy has been shown to reduce proteinuria and delay progression to end-stage renal disease in IgAN[42, 43], these EV-associated metabolic changes likely reflect endogenous steroid-related metabolic processes rather than the direct effects of exogenous steroid treatment. The relationship between endogenous steroid metabolism captured by EVs and clinical responsiveness to steroid therapy therefore warrants further investigation.

Several limitations of this study should be acknowledged. First, the relatively small sample size and single-center design may limit the generalizability of our findings and reduce statistical power to detect subtle proteomic and metabolomic differences. Second, although urinary EV proteomics provides valuable insights for biomarker discovery, the limited number of DEPs restricts comprehensive pathway analysis and precludes causal inference regarding disease mechanisms. Finally, due to the lack of authentic reference standards and confirmatory MS/MS spectra, metabolite identification could only be assigned confidence level 3[44]. Future studies involving larger, multicenter cohorts and mechanistic investigations are required to validate and extend these findings.

## Conclusion

As the first comprehensive study investigating plasma- and urinary EV-based proteomic and metabolomic biomarkers for IgAN, our findings provide strong evidence supporting the utility of plasma- and urinary EV proteomics and metabolomics for early biomarker discovery. Venn diagram analysis identified 25 candidate DEPs and 1546 candidate DEFs with potential for early detection, among which C4BPA, PROS1, and metabolite features with formulas C27H44O, C30H50O, and C28H46O showed particular promise as indicators of IgAN disease status and progression.

## List of abbreviations

AUC: Area under the curve
CKD: Chronic kidney disease
C4BPA: C4b-binding protein alpha chain
C4BPB: C4b-binding protein beta chain
DEFs: Differentially expressed features
DEPs: Differentially expressed proteins
EVs: Extracellular vesicles
GC: Vitamin D–binding protein
HBB: Hemoglobin subunit beta
IgAN: Immunoglobulin A nephropathy
NTA: Nanoparticle tracking analysis
pEVs: Plasma extracellular vesicles
PROS1: Vitamin K–dependent protein S
ROC: Receiver operating characteristic
sPLS-DA: Sparse partial least squares-discriminant analysis
TEM: Transmission electron microscopy
uEVs: Urinary extracellular vesicles

## Declarations

### Ethics approval and consent to participate

This study was approved by the Joint Institutional Review process of the Taipei Medical University – Joint Institutional Review Board No. N201704064). Signed informed consent was obtained from all participants recruited in the study.

### Consent for publication

Not applicable

### Availability of data and materials

All data related to this article are disclosed or available upon request from the corresponding author.

### Authors’ contributions

Conceptualization: YHL, ILT, CCK; Investigation: YHL, TYC; Formal analysis: YHL, TYC, JYP; Visualization: YHL, TYC, ILT; Validation: YHL, TYC; Resources: CCK; Writing—Original Draft: YHL, TYC, ILT; Writing—Review & Editing: ILT, CCK; Supervision: ILT, CCK; Project administration: ILT, CCK; Funding acquisition: ILT, CCK.

### Data statement

Most of the data and information supporting the findings of this study are available within the main text. The raw mass spectrometry proteomics data related to Figures 2 and 3 have been deposited in the ProteomeXchange Consortium via the PRIDE partner repository under the dataset identifier PXD074396. Additional datasets are available from the corresponding author, CK, on reasonable request.

## Acknowledgments

We thank the mass spectrometry technical research services provided by Consortia of Key Technologies and Instrumentation Center, National Taiwan University.

