## Supplementary figures and images for "Multi-Omics Characterization of Plasma and Urine Extracellular Vesicles Identifies Non-Invasive Biomarkers for IgA Nephropathy"

### Figure S1

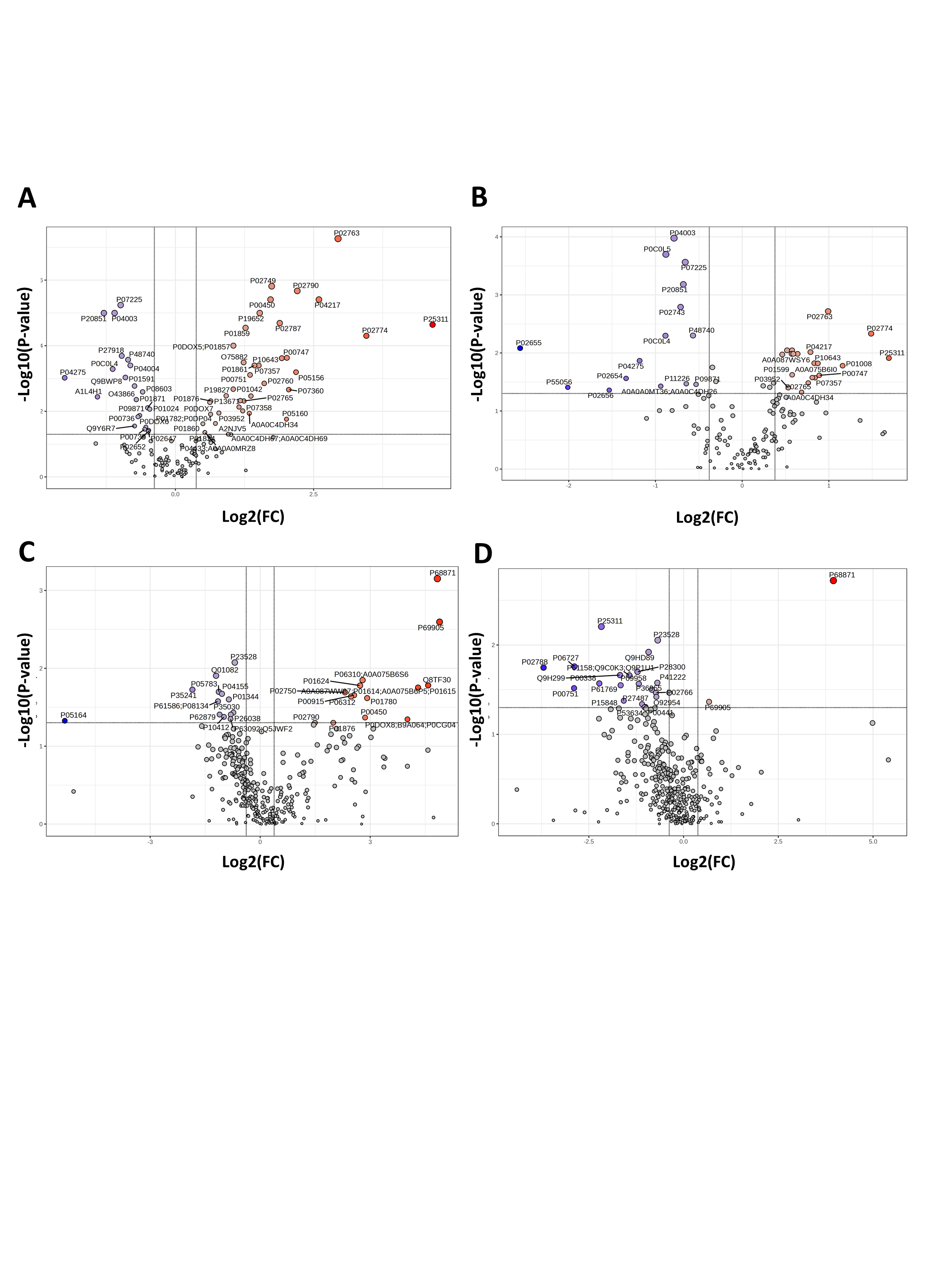

### Figure S2

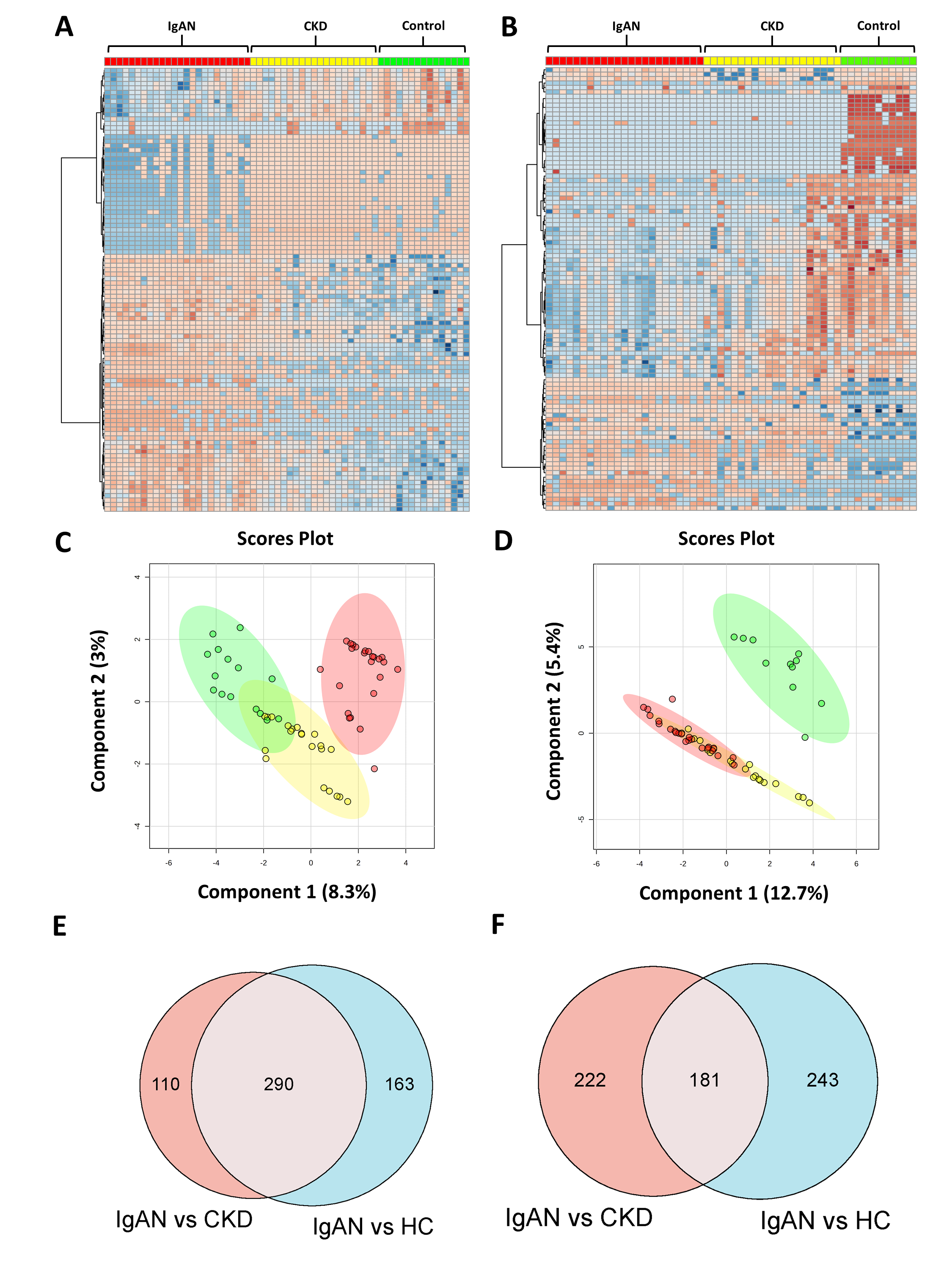

### Figure S3

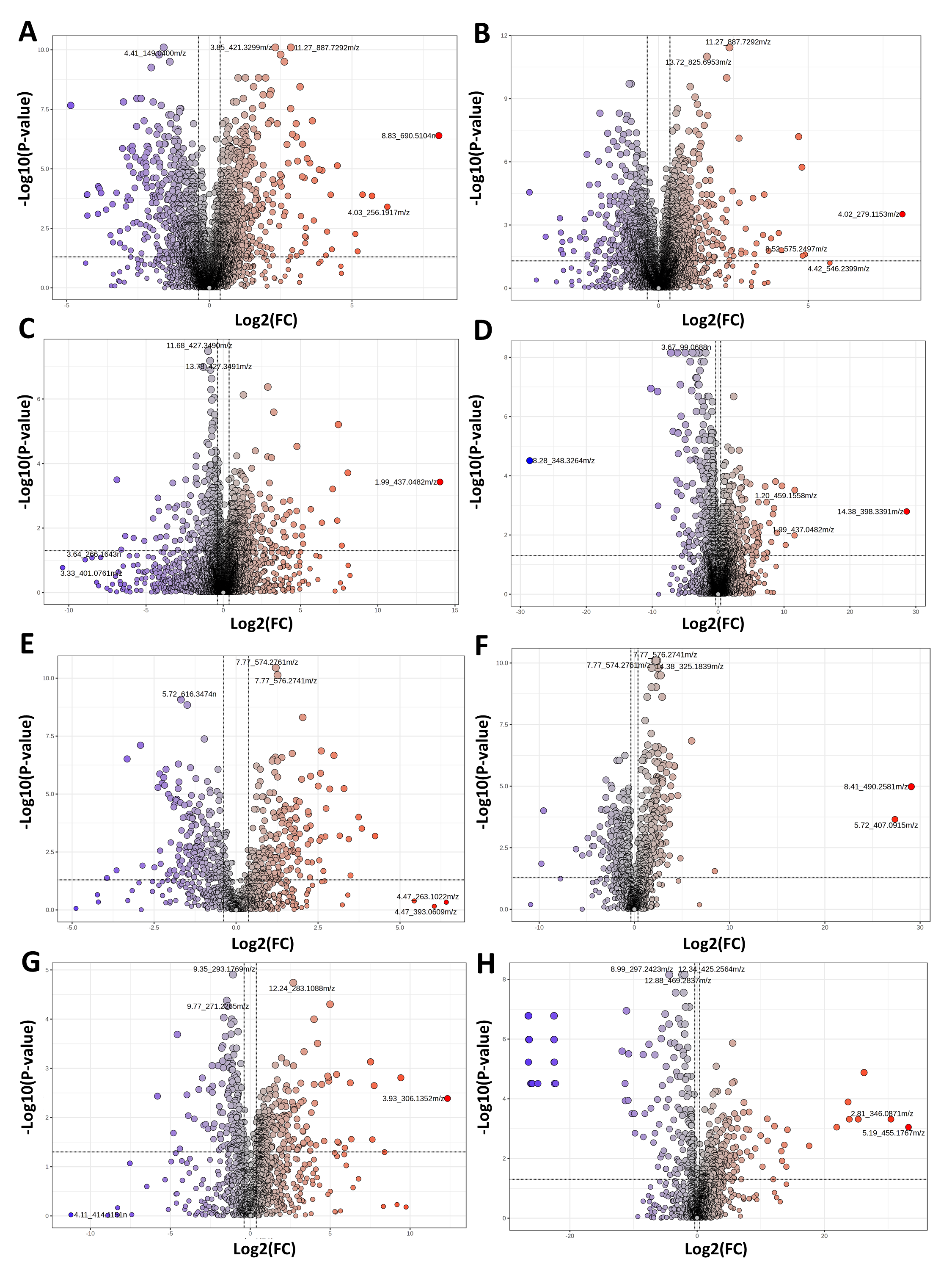

### Figure S4

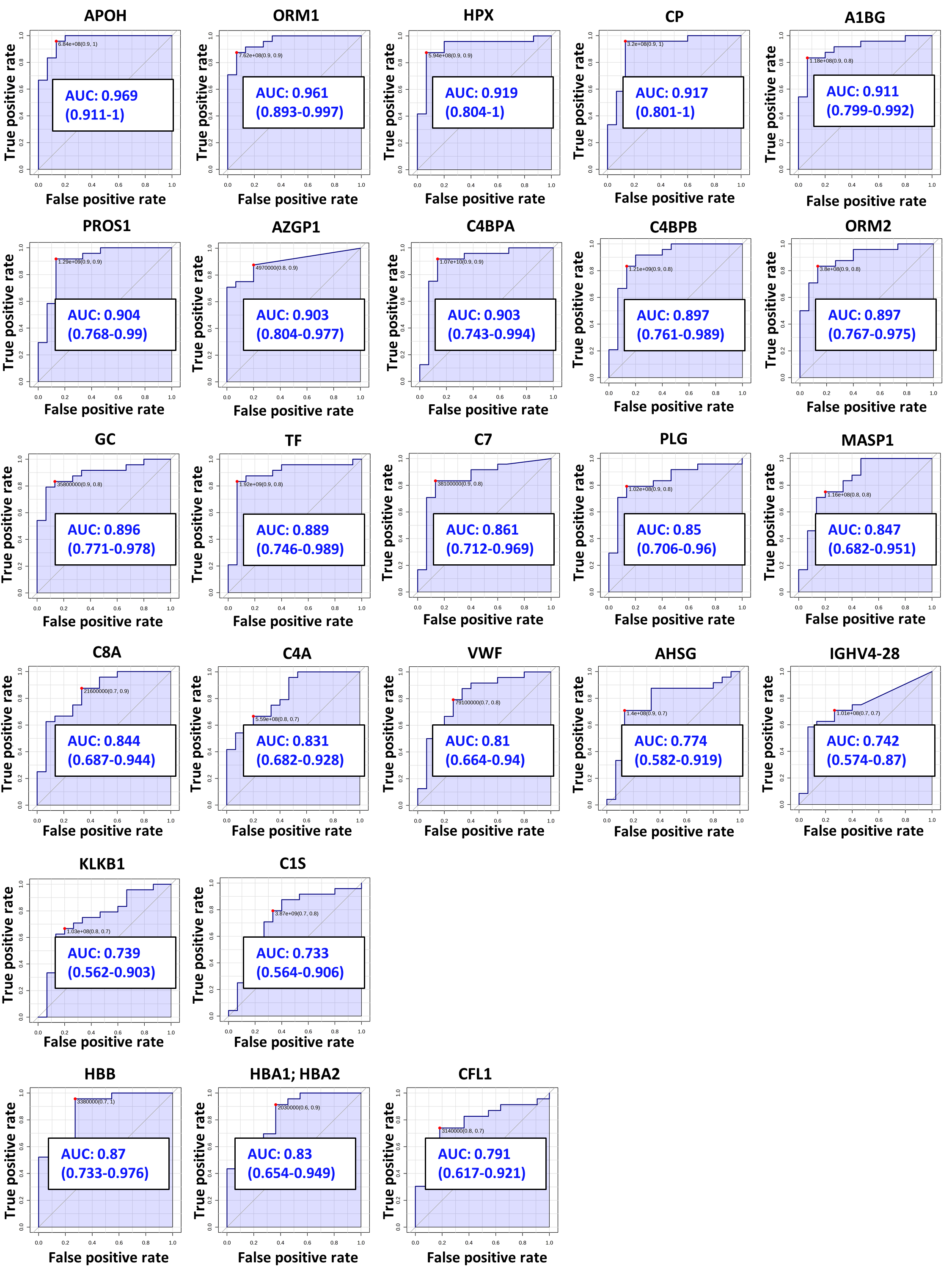
